# Elucidating the differences in the molecular mechanism of receptor binding between 2019-nCoV and the SARS-CoV viruses using computational tools

**DOI:** 10.1101/2020.04.21.053009

**Authors:** Hien T. T. Lai, Ly H. Nguyen, Agata Kranjc, Toan T. Nguyen, Duc Nguyen-Manh

## Abstract

The outbreak of the 2019-nCoV coronavirus causing severe acute respiratory syndrome which can be fatal, especially in elderly population, has been declared a pandemic by the World Health Organization. Many biotechnology laboratories are rushing to develop therapeutic antibodies and antiviral drugs for treatment of this viral disease. The viral CoV spike (S) glycoprotein is one of the main targets for pharmacological intervention. Its receptor-binding domain (RBD) interacts with the human ACE2 receptor ensuring the entry of the viral genomes into the host cell. In this work, we report on the differences in the binding of the RBD of the previous coronavirus SARS-CoV and of the newer 2019-nCoV coronavirus to the human ACE2 receptor using atomistic molecular dynamics techniques. Our results show major mutations in the 2019-nCoV RBD with respect to the SARS-CoV RBD occurring at the interface of RBD-ACE2 complex. These mutations make the 2019-nCoV RBD protein backbone much more flexible, hydrophobic interactions are reduced and additional polar/charged residues appear at the interface. We observe that higher flexibility of the 2019-nCoV RBD with respect to the SARS-CoV RBD leads to a bigger binding interface between the 2019-nCoV RBD and ACE2 and to about 20% more contacts between them in comparison with SARS-CoV. Taken together, the 2019-nCoV RBD shows more stable binding interface and higher binding affinity for the ACE2 receptor. The mutations not only stabilize the binding interface, they also lead to overall more stable 2019-nCoV RBD protein structure, even far from the binding interface. Our results on the molecular differences in the binding between the two viruses can provide important inputs for development of appropriate antiviral treatments of the new viruses, addressing the necessity of ongoing pandemics.

## Introduction

The COVID-19 disease that has rapidly spread from the Chinese Wuhan city to the rest of the world is caused by the 2019 novel coronavirus (2019-nCoV) named also the severe acute respiratory syndrome corona virus 2 (SARS-CoV-2).^1^ The sequencing of its genome revealed that it is closely related to other coronaviruses, such as severe acute respiratory syndrome coronavirus (SARS-CoV) and Middle East respiratory syndrome coronavirus (MERS-CoV), which have killed hundreds of people in the last two decades.^2,3^

Coronaviruses are enveloped, positive sense, single-stranded RNA viruses that carry on their surface spike-like projections giving it a crown-like appearance under the electron microscope; hence the name coronavirus.^4^ These spikes (S) enable the fusion between viral and host membranes and are essential for the beginning of the enveloped virus infection.^5,6^ They are composed of a large ectodomain, a trans-membrane anchor and a short intracellular tail (Fig. 1). The ectodomain consists of a receptor-binding subunit S1 and a membrane-fusion subunit S2 which are crucial for binding of the virus to the host cell surface and for entry of the viral genomes into the target cell, respectively.^7–11^ The S1 subunit is composed of N-terminal domain (S1-NTD) and C-terminal domain (S1-CTD), which can both function as the Receptor Binding Domain (RBD), recognizing and binding to the host protein receptor. The cellular entry receptor for the SARS-CoV is Angiotensin Converting Enzyme 2 (ACE2).^12^ The RBD of SARS-CoV and SARS-CoV-2 share significant sequence similarity, therefore several research groups investigated if SARS-CoV-2 uses the same cellular entry receptor.^2,13–16^ And they all confirmed that this is indeed the case.

**Figure 1:**
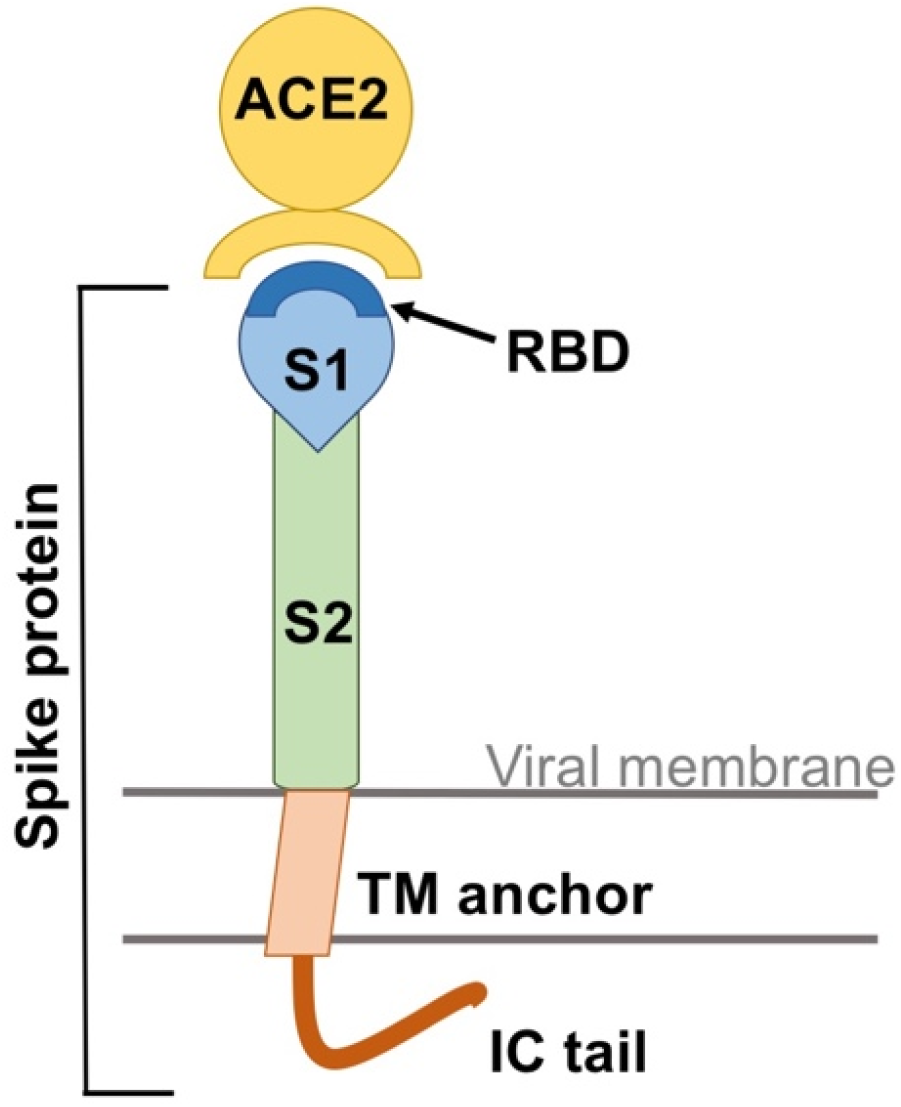
A schematic view of the coronavirus spike protein. The spike protein bears a large ectodomain that is composed of a subunit S1 which has at the top the Receptor Binding Domain (RBD) and the subunit S2. It further features the transmembrane anchor (TM anchor) and the intracellular tail (IC tail). ACE2 is the Angiotensin Converting Enzyme 2, the main host entry receptor for SARS-CoV and SARS-CoV-2.

The ACE2 enzyme is a part of the renin-angiotensin-aldosterone system (RAAS) that controls blood pressure, fluid and electrolyte balance, systemic vascular resistance, tissue damage etc.^17^ The primary physiological role of the ACE2 is to convert a peptide hormone angiotensin II into angiotensin (1-7), that have protective role as they diminish the inflammation and fibrosis.^17,18^ The SARS-CoV and SARS-CoV-2 viruses bind to ACE2 preventing the metabolism of the angiotensin II, which accumulates and can cause deleterious effects. There are many controversial debates about the beneficial or harmful role of the RAAS inhibitors in patients with COVID-19.^18–22^ Number of experts defend their beneficial role, but the qualified clinical studies will have to be done to answer this important question. The SARS-CoV and SARS-CoV-2 are closely related, nonetheless, significant differences were reported for their protein structures, importantly the mutations in the SARS-CoV-2 RBD domain with respect to the SARS-CoV that can impact the binding affinity for the host receptor.^14,23^ In our study we aimed to compare the structural and energetic differences in the binding of the RBDs of the SARS-CoV-2 and SARS-CoV to the ACE2 receptor by Molecular Dynamics (MD) simulations. The viral RBD represents one of the main possible targets for the development of antiviral drugs, therefore, understanding the binding between the viral RBD and ACE2 can be very useful for rational drug design. We exploited the fact that different 3D protein structures of the ACE2 in complex with the RBD of SARS-CoV and SARS-CoV-2 are available in the Protein Data Bank: (i) SARS-CoV RBD in complex with ACE2, PDB code: 2AJF^24^ (ii) SARS-CoV-2 RBD in complex with ACE2, PDB code: 6VW1^25^ (iii) SARS-CoV-2 RBD in complex with ACE2, PDB code: 6M0J.^26^ By using two SARS-CoV-2 samples and one SARS-CoV sample, we can effectively compare the variations among the viruses, as well as identify the important and unimportant mutations and interactions.

Molecular dynamics is performed to investigate the characteristics of the binding mechanism between viral RBDs and its receptor. Sequence, structural and dynamical analyses are carried out to identify similarities and differences between the new SARS-CoV-2 viruses and the previous SARS-CoV virus. Our results provide a molecular understanding of the binding complex and show that the new viruses bind more strongly to the receptor. Despite variations between the sequences of the two SARS-CoV-2 viruses, the two samples of the new viruses behave very similarly, and have similar binding properties for the human receptor. By eliminating unimportant differences in the sequences of the new viruses, and comparing them to the sequence of the old SARS-CoV virus, four important mutations between SARS-CoV-2 and SARS-CoV are identified: Y to L455, VP to EI471-472, -PP to GVE482-484, and Y to Q498 (the indices are that of SARS-CoV-2 virus, which is shifted higher by 13 compared to SARS-CoV virus numbering). They are shown structurally to be at or close to the binding interface with the receptor protein. The mutations lead to higher flexibility of the backbones by substitution of Prolines and insertion of Glycine. The mutations also reduce number of hydrophobic interactions at the interface since hydrophobic residues are substituted with charged or polar residues. With higher backbone flexibility, new residues can move closer to the receptor interface, increasing the number of non-bonded interactions between the proteins. Specifically, the mutations eliminate one hydrogen bonding with the receptor, but increase the coordination number (number of contacts) by 20%. This leads to stronger non-bonded interactions (electrostatics and van der Waals) with the receptor compared to the SARS-CoV virus. The overall results suggest that the binding energy in the new SARS-CoV-2 viruses is slightly higher relative to the SARS-CoV virus. This is in reasonable agreement with experimental results reporting that the new corona virus has 10-20 times higher binding affinity for the ACE2 receptor than the old SARS-CoV.^27^ Structurally, stronger binding is shown to not only stabilize the binding interface, but also to stabilize the overall structural fold of the viral RBD of the new SARS-CoV-2 viruses.

This paper is organized as follows. After the introduction in Section 1, the detail of the computational procedure is presented in Section 2. In Section 3, the results are presented and discussed. We conclude in Section 4.

## Materials and Methods

The RBD - ACE2 complexes were obtained from the RCSB PDB database with ID: 6VW1,^25^ 6M0J26 for the new SARS-CoV-2 viruses, and 2AJF24 for the SARS-CoV virus. For easy identification, these systems are named 6VW1, 6M0J and 2AJF respectively. The 6VW1 and 2AJF systems contain two complexes, chains A and E, chains B and F. The 6M0J contains only one complex, chains A and E. We keep only the complex of chain A (ACE2 receptor) with chain E (viral RBD) in these samples to better compare them to one another. This is also understandable since the complex of chain B and F is a near 180° rotation symmetry of the complex of chain A and E. It is an artifact of protein crystallization, not actual arrangement of the complexes in real biological system.

There are some notable structural elements in these complexes that need special care when setting up the simulation systems such as: the disulfide bonds between various pairs of Cysteine residues; the N–linked glycosylations of various Asparagine residues; and the ZnGlu_2_His_2_ zinc–finger structure that needs proper protonation states of the associated amino acids.^28^ Details of these structural elements are listed in Table 1 for better identification in later discussion of the simulation results..

**Table 1:** Detail information of various special structural elements of the SARS-CoVs RBD - ACE2 complexes that need proper care when setting up the system.

| Models |  | S-S bridges | Zinc finger | Glycosylation |
| --- | --- | --- | --- | --- |
| 6VW1 | ACE2 | C133 - C141<br>C344 - C361<br>C530 - C542 | H374<br>E375<br>H378<br>E402 | bDMan(1→4)bDGlcNAc(1→4)bDGlcNAc(1→)N90<br>bDMan(1→4)bDGlcNAc(1→4)bDGlcNAc(1→)N546<br>bDGlcNAc(1→4)bDGlcNAc(1→)N53<br>bDGlcNAc(1→4)bDGlcNAc(1→)N322<br>bDGlcNAc(1→)N103 |
|  | RBD | C336 - C361<br>C379 - C432<br>C391 - C525<br>C480 - C488 |  | bDMan(1→4)bDGlcNAc(1→4)bDGlcNAc(1→)N343 |
| 6M0J | ACE2 | C133 - C141<br>C344 - C361<br>C530 - C542 | H374<br>E375<br>H378<br>E402 | bDGlcNAc(1→)N90<br>bDGlcNAc(1→)N546<br>bDGlcNAc(1→)N322 |
|  | RBD | C336 - C361<br>C379 - C432<br>C391 - C525<br>C480 - C488 |  | bDGlcNAc(1→)N343 |
| 2AJF | ACE2 | C133 - C141<br>C344 - C361<br>C530 - C542 | H374<br>E375<br>H378<br>E402 | bDMan(1→4)bDGlcNAc(1→4)bDGlcNAc(1→)N90<br>bDGlcNAc(1→)N53<br>bDGlcNAc(1→)N322<br>bDGlcNAc(1→)N546 |
|  | RBD | C323 - C348<br>C366 - C419<br>C467 - C474 |  | bDGlcNAc(1→)N343 |

For alignment of the primary sequences of the proteins, ClustalW web-server^29^ with BLO-SUM matrix,^30^ BioEdit software^31^ are used. Missing residues in the central protein structure were added using homology modeling method.^32–34^ The initial systems for simulation were created using CHARMM-GUI web-server^35^ and then modified manually to properly describe some of above structural elements. Molecular Dynamics (MD) simulations^36^ are performed on the systems using the GROMACS/2018.6 software package.^37^ Charmm-36 force-field^38^ was chosen for parametrization of the proteins and ions. GLYCAM06 force-field^39^ was used to parametrize the glycans and TIP3P^40^ model is used for water in the explicit solvent simulation model. After solvation, sodium and chlorine ions are added to the system to neutralize the total charges and to set physiological electrolyte concentration of solution at 150 mM NaCl. The simulation box size was chosen so that the proteins in neighbor periodic box are at least 3 nm apart from each other. Since the electrostatic screening length at 150mM NaCl concentration is about 7Å, this 3 nm distance is more than enough to eliminate the finite size effect due to long range electrostatic interactions among proteins in neighboring simulation boxes, yet small enough to keep the size of the system manageable with our current computational resources. The total numbers of molecules, residues and atoms for the three systems simulated are listed in Table 2: The temperature of 310 K and the pressure of 1 atm are maintained by the Nose-Hoover thermostat and the Parrinello-Rahman barostat. The Particle Mesh Ewald (PME) method is used to treat the long-range electrostatic interaction with a real space cutoff of 1.2 nm. Van der Waals interactions are also cut off at 1.2 nm, with the appropriate cut-off corrections added to pressure and energy. All hydrogen bonds are constrained by the LINCS method.^41^ The systems are equilibrated in NPT ensemble for 150 ps at timestep of 1 fs. After that, 350 ns MD production run at timestep of 2 fs is performed for statistics.

**Table 2:** The molecules simulated for each systems.

|  | **6VW1** |  | **6M0J** |  | **2AJF** |  |
| --- | --- | --- | --- | --- | --- | --- |
|  | # Residues | # Atoms | # Residues | # Atoms | # Residues | # Atoms |
| ACE2 | 597 | 9802 | 597 | 9598 | 597 | 9673 |
| RBD | 194 | 3070 | 194 | 3020 | 180 | 2848 |
| H<sub>2</sub>O | 87382 | 262146 | 76599 | 229797 | 66380 | 199140 |
| Na<sup>+</sup> |  | 275 |  | 241 |  | 213 |
| Cl<sup>-</sup> |  | 250 |  | 218 |  | 189 |
| Zn<sup>2+</sup> |  | 1 |  | 1 |  | 1 |
| Total |  | 275544 |  | 242874 |  | 212064 |

Standard analyses such as mean deviation, mean fluctuation, hydrogen bonding are performed using GROMACS software. Visual Molecular Dynamics (VMD) program^42^ is used for visualization, spatial inspection and some surface analyses. Some collective variable analyses are performed using PLUMED software.^43^ In-house Python scripts are used to integrate various tools for Principal Component Analysis (PCA), setting up the systems, process simulation data, graphical plottings. SciPy python package is used for calculating probability density, Gaussian quadrature from which the difference in the binding free energy of the proteins is estimated.

## Results and discussions

### Preliminary sequence and structural analyses

Before going into detailed discussion of simulation results, let us analyse the sequences and structures of the three RBD–ACE2 complexes of the experimental samples, 2AJF for SARS-CoV virus, 6VW1 and 6M0J for SARS-CoV-2 viruses. This helps to clarify common features as well as differences between SARS-CoV and SARS-CoV-2 viruses. This information will be valuable for later interpretation and discussion of results of our MD simulations.

#### The human ACE2 receptor

In the experimental X–ray crystal structures of 2AJF,^24^ 6VW1^25^ and 6M0J,^26^ sequences of the human ACE2 receptor are identical but glycosylations of the human ACE2 protein in these complexes show some small variations (see Table 1). For the 2AJF sample, the N-link glycosylations occur at the four Asparagine residues, N53, N90, N322, and N546. They link to chains of two or three N-Acetyl-D-Glucosamine (NAG) and may end with *β*-D-Mannose (BMA) molecules. For the 6VW1 sample, the same residues are also N-link glycosylated, albeit with slightly different length of the oligosacharides. Additionally, residue N103 is also glycosylated with a single NAG residue. For the 6M0J sample, all three Asparagine residues, N90, N322, and N546 are N–link glycosylated with a single NAG sugar molecule.

The N–link glycosylation plays an important role in the protein structure stability. Despite different number of glycosylated sites in studied ACE2 proteins, we will see later in our MD simulation results that the three ACE2 proteins have nearly identical structural and dynamical properties in low energy modes. Only the 6M0J sample with less glycosylation degrees shows some differences at localized high energy dynamical modes, but these differences do not influence the binding interface of the viral RBD and the ACE2 receptor.

#### The viral receptor-binding domain

Unlike the human receptor, the sequences of viruses’ receptor-binding domain (RBD) of the coronaviruses in all three structures, 2AJF, 6VW1 and 6M0J are different to varying degrees of conservations. In Fig. 2, these three sequences of the viral RBD are aligned using ClustalW web-server.^29^ The sequence identity between the two SARS-CoV-2 coronaviruses is 85%. The sequence identity between SARS-CoV (2AJF) with two SARS-CoV-2 samples (6M0J and 6VW1) are 72% and 85%, respectively. Thus, these RBD of coronaviruses are very similar in sequence and structure. This is expected because they are closely related in the coronavirus family.

**Figure 2:**
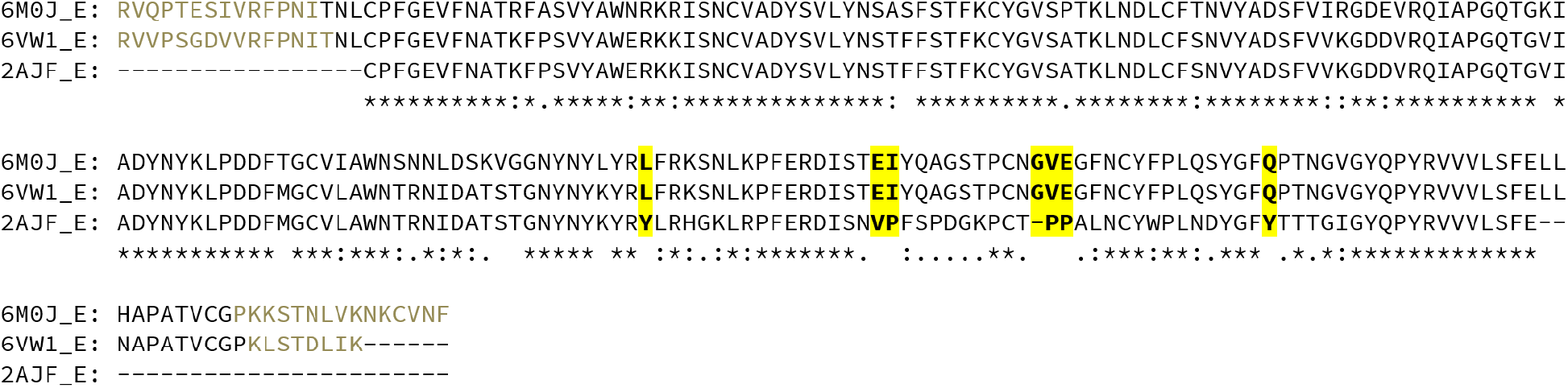
Sequence alignments of the viral RBD of two SARS-CoV-2 stuctures (6M0J and 6VW1) and the SARS-CoV structure (2AJF). The *, :, or . notations below the sequence denote a fully, strongly, or weaker conserved mutation respectively. A blank space below the sequences means a non-conserved mutation. Different mutations emerge between the compared RBDs, but four groups of mutations (highlighted in yellow) show aminoacids fully conserved between SARS-CoV-2 samples and non-conserved in the SARS-CoV sequence (see text for more discussion). The grayed–out residues at the end of the sequences are flexible N- and C-termini loops that are missing in the PDB crystal structure.

Since we are looking for major differences between SARS-CoV and SARS-CoV-2, we only look for aminoacids that are the *same in the two SARS-CoV-2* viruses but are *not-conserved* with respect to SARS-CoV virus. Using these criteria, we identify four locations of important mutations. These are highlighted in yellow in Fig. 2. Remarkably, upon spatial inspection, all these four locations are at or close to the binding interface with the human ACE2 receptor. This is a strong hint that these mutations can be critical for the stronger receptor binding of the new viruses. We will focus more on these mutations in later discussions of results of MD analyses.

Before discussing the MD results, a quick scan on the physical changes at some of these mutations can already be made:

- **Y** to **L455**: this mutation in SARS-CoV-2 leads to the loss of the aromatic ring and especially the hydroxyl group of Tyrosine amino acid. So it removes the possibility for hydrogen bond with the receptor counterpart residue Glu35. In MD analyses, we will see that the loss of hydrogen bond binding at this location is compensated with other non-bonded interactions.
- **VP** mutated to **EI** at index 471-472 of SARS-CoV-2: this mutation substitutes the rigid Proline residue and adds the negatively charged Glutamic Acid E471. This increases the flexibility of the protein backbone.
- **-PP** to **GVE** at index 482-484 of SARS-CoV-2: The insertion of Glycine and the substitutions of two Proline amino acids clearly make this segment much more flexible. This flexibility allows the segment to move closer to its receptor and enables the negatively charged E484 residue to form electrostatic bond with the positively charged residue K31 of the human ACE2 receptor.
- **Y** to **Q498** mutation in SARS-CoV-2 turns the T-stacking hydrophobic interaction between the two tyrosines at the RBD-ACE2 interface into electrostatic interactions with the opposing Q42 residue of the ACE2 receptor.

In summary, these mutations increase the flexibility of the backbone atoms at the viral binding interface and they add to this interface polar or charged residues. Higher flexibility allows them to adapt better to the receptor binding interface, increasing the number of contacts between the two proteins. The increase in the non-bonded electrostatic and van der Waals interactions then leads to higher binding affinity as found later by MD simulation.

### Residues at N- and C-termini of SARS-CoV-2 viral RBD

Fig. 2 shows, the viral RBD of SARS-CoV-2 samples have some additional residues at the N- and C-termini compared to the SARS-CoV sample. However, Fig. 3 shows that these RBD residues are located far away from the binding interface with the human ACE2 receptor. Because our main goal is to discern the differences between SARS-CoV and SARS-CoV-2 receptor binding, whenever appropriate when aligning the structures for analyses, we use *only the common “core”* part of the three viral RBD proteins (from amino acid sequence CPFGE to VLSFE in Fig. 2).

Other differences among the viral RBD structures are the missing residues at the N- and C-termini. In Fig. 2, the grayed–out residues at the two ends of SARS-CoV-2 sequences are missing in the protein experimental crystal structures since the precise 3D structures of the termini loops cannot be determined by crystallography because of their flexibility. To find out if these residues may play an important role for the stability of the structure, we simulate two different 6VW1 systems, the first with original PDB structure and the second where the missing termini residues are added to the PDB structure by homology modelling.^32–34^ Both systems are simulated for 200 ns and statistics are collected from 50 ns onward. By comparing to the systems with and without added missing residues, we see no noticeable differences in the structures of the binding interface. In Fig. 4, the mean fluctuations of C_*α*_ atoms in the core region of the viral RBD are plotted for the systems with or without terminal residues added (blue and red curves, respectively). As one can see, the difference among the systems is minimal, mostly less than 0.2Å.

**Figure 3:**
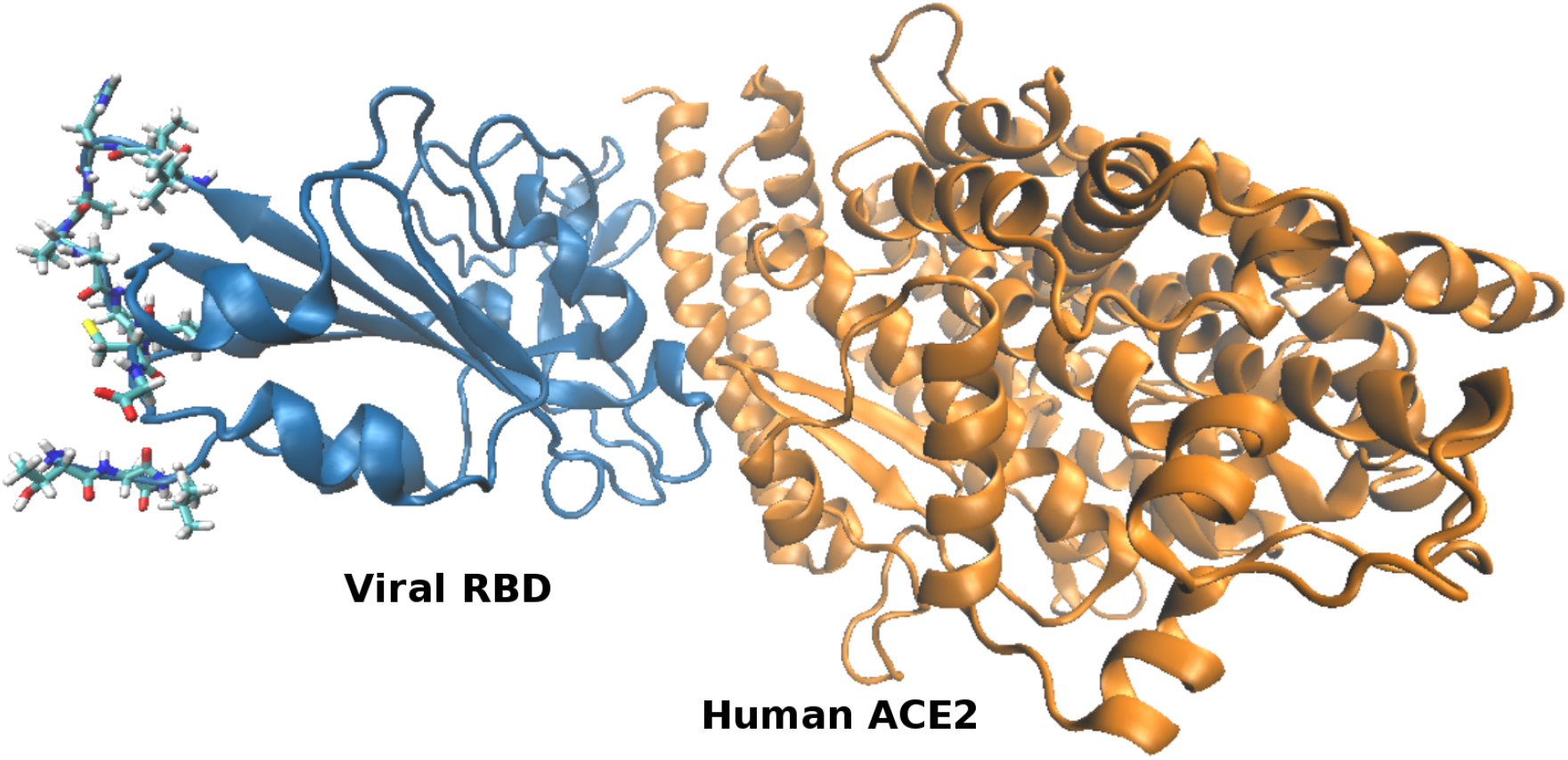
Compared to the experimentally-resolved viral RBD structure of SARS-CoV, the SARS-CoV-2 viral RBD structure has additional residues at the N- and C-terminals. However, these residues (shown in sticks) are far from the binding interface and will not be used for alignment of structures. In this figure, the 6VW1 sample is shown. The 6M0J structure is very similar.

**Figure 4:**
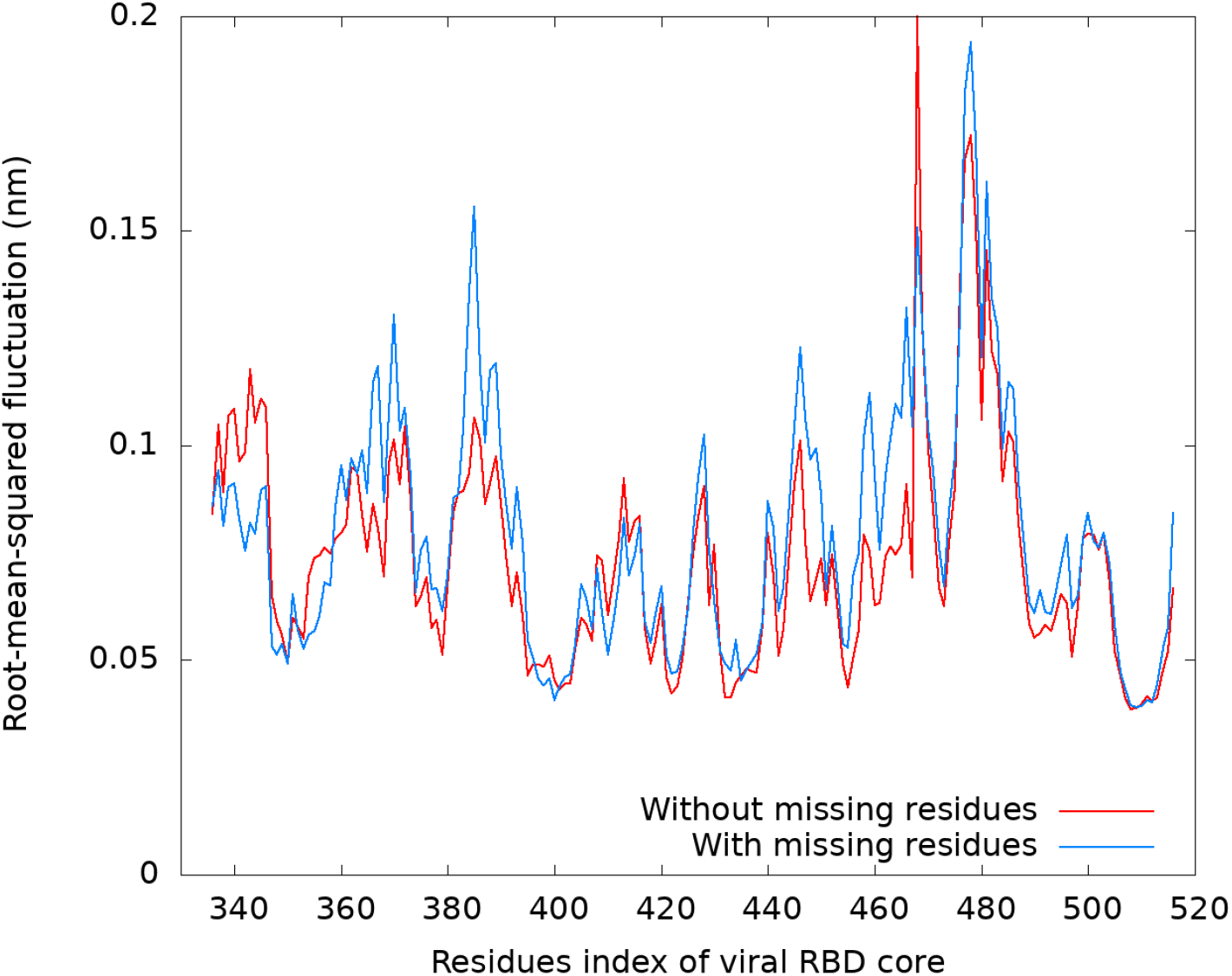
Root-mean-square fluctuations of the “core” region of the viral RBD structure of the 6VW1 complex. The blue curve is for the complex where missing residues at the termini are added by homology modeling method. The red curve is for the original complex.

Quantities such as radius of gyrations, moment of inertia along principle axes of systems with and without missing residues also show identical values within standard uncertainty as shown in Table 3. Thus, one can conclude that adding missing terminal residues does not alter significantly the structure of the core binding region of the viral RBD. On the other hand, since these flexible and disordered terminal loops extend far into the aqueous solution, they significantly increase the size of the simulation box. As a result from these considerations, for the rest of this work, we simulate only the residues present in the crystal structure without the missing residues at the termini to save computational resources.

**Table 3:** Various physical quantities of the core region of viral RBD for the 6VW1 sample show no noticeable differences with or without missing terminal residues added.

|  | With missing residues | Without missing residues |
| --- | --- | --- |
| Total moment of inertia, $I$ (amu nm<sup>2</sup>) | $78791 \pm 806$ | $78540 \pm 940$ |
| Moment of inertia, $I_1$ (amu nm<sup>2</sup>) | $24428 \pm 320$ | $24211 \pm 272$ |
| Moment of inertia, $I_2$ (amu nm<sup>2</sup>) | $48155 \pm 678$ | $48231 \pm 717$ |
| Moment of inertia, $I_3$ (amu nm<sup>2</sup>) | $57376 \pm 624$ | $57060 \pm 749$ |
| Radius of gyration, $R_g$ (nm) | $1.781 \pm 0.008$ | $1.778 \pm 0.010$ |
| Radius of gyration, $R_{gx}$ (nm) | $1.494 \pm 0.091$ | $1.516 \pm 0.100$ |
| Radius of gyration, $R_{gy}$ (nm) | $1.531 \pm 0.110$ | $1.512 \pm 0.086$ |
| Radius of gyration, $R_{gz}$ (nm) | $1.317 \pm 0.115$ | $1.304 \pm 0.148$ |

The ACE2 protein of the 6VW1 system misses a D615 residue at the C–terminal compared to the ACE2 protein in the 2AJF and 6M0J systems. Since this is just one residue, costing minimal computing resource, we add this missing residue in the simulated systems. The three systems simulated thus have identical ACE2 protein sequences. Most of the differences are due to changes in the viral RBD sequences.

### Deviations and fluctuations of the structural backbone atoms

Let us now present and discuss the results of our molecular dynamics (MD) simulations of the three systems and compare the differences between SARS-CoV and SARS-CoV-2 samples. As a standard procedure, one first looks at the deviations of the structural proteins from their native experimental structures. In Fig. 5, the root-mean-square deviations (RMSD) of backbone atoms in the three systems are plotted as function of time. Fig. 5(a) is plotted for the human ACE2 receptor, while Fig. 5(b) is plotted for the viral RBD protein. The viral RBD, being smaller (~ 200 residues) shows faster saturation time (at around 50ns) than the ACE2 receptor (~ 600 residues) whose saturation time is at around 100 ns or longer. Among SARS-CoV versus SARS-CoV-2 viruses, the viral RBD is remarkably more stable in the new viruses as they both deviate only 1.5Å from the native structure while that of the old SARS-CoV virus shows about 3Å deviation. Similarly to the RBD, the human ACE2 in the SARS-CoV-2 systems are more stable and show lower RMSD values than in the SARS-CoV complex. The ACE2 receptor in 2AJF system also indicates longer relaxation time. Nevertheless, both proteins in all systems are stable with RSMD less than 4Å deviation from native structure. As a result of this RMSD analysis, in all subsequent equilibirium statistical analyses, the first 100 ns of the MD trajectories will be dropped.

**Figure 5:**
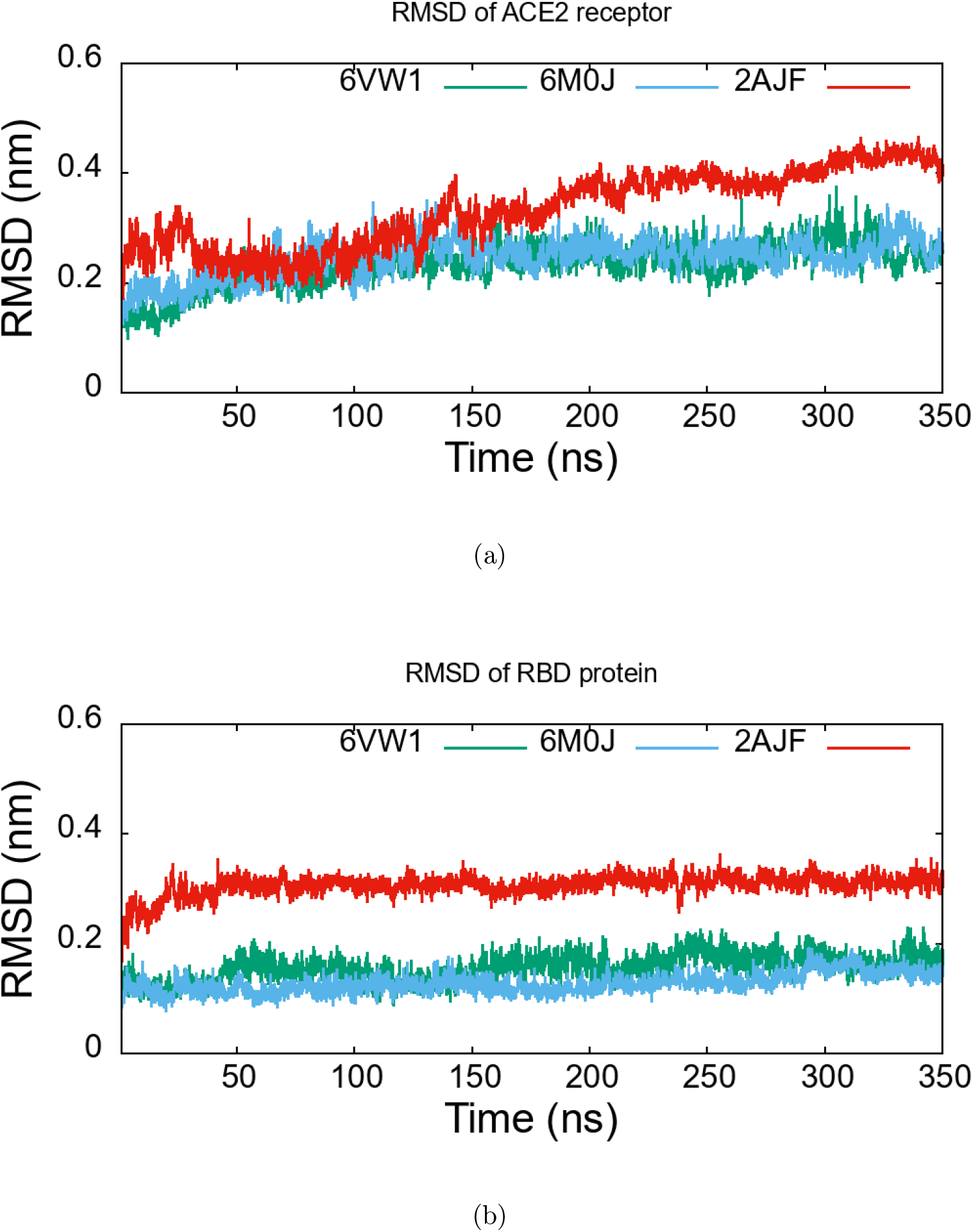
The root-mean-square deviation of backbone atoms (backbone RMSD) of the human ACE2 receptor (a) and the viral RBD protein (b) in MD simulations from the initial crystal structure. Both systems of SARS-CoV-2 (green and cyan curves) show smaller deviations with faster time to saturation than that of SARS-CoV system (red curve). Both proteins are stable with deviations less than 4Å from the native structure.

Next, let us look at the root mean square fluctuations (RMSF) of the atoms. This fluctuation is due to thermal effect of finite temperature (at 310 K) compared to the low temperature of experimental crystallized structure where proteins are frozen. It is a good measurement of the backbone stability of the proteins. In Fig. 6(a), RMSF of the backbone C_*α*_ atoms of the human ACE2 receptor in the three systems are plotted for individual residues. One can see from this figure that this receptor behaves similarly across the three systems, with variations less than 0.5Å from one system to another. Our MD simulations show as well that the different ACE2 glycosylation states (see Table 1 for more details) have minimal influence on the rigidity of the protein backbones. One noteworthy difference is the slightly higher fluctuations of ACE2 residues 134-177, 262-278, 489-500 in the 6M0J system compared to the other two. As it turns out, these residues surround a Cl^−^ ion in the 2AJF and 6VW1 systems (see Fig. 6(b)). For unknown reasons, this chlorine ion is absent in the experimental crystal structure of 6M0J system. These residues are at the other extreme of the ACE2 receptor, far from the binding interface, so we believe that the slightly higher fluctuations in this region have no important consequences.

**Figure 6:**
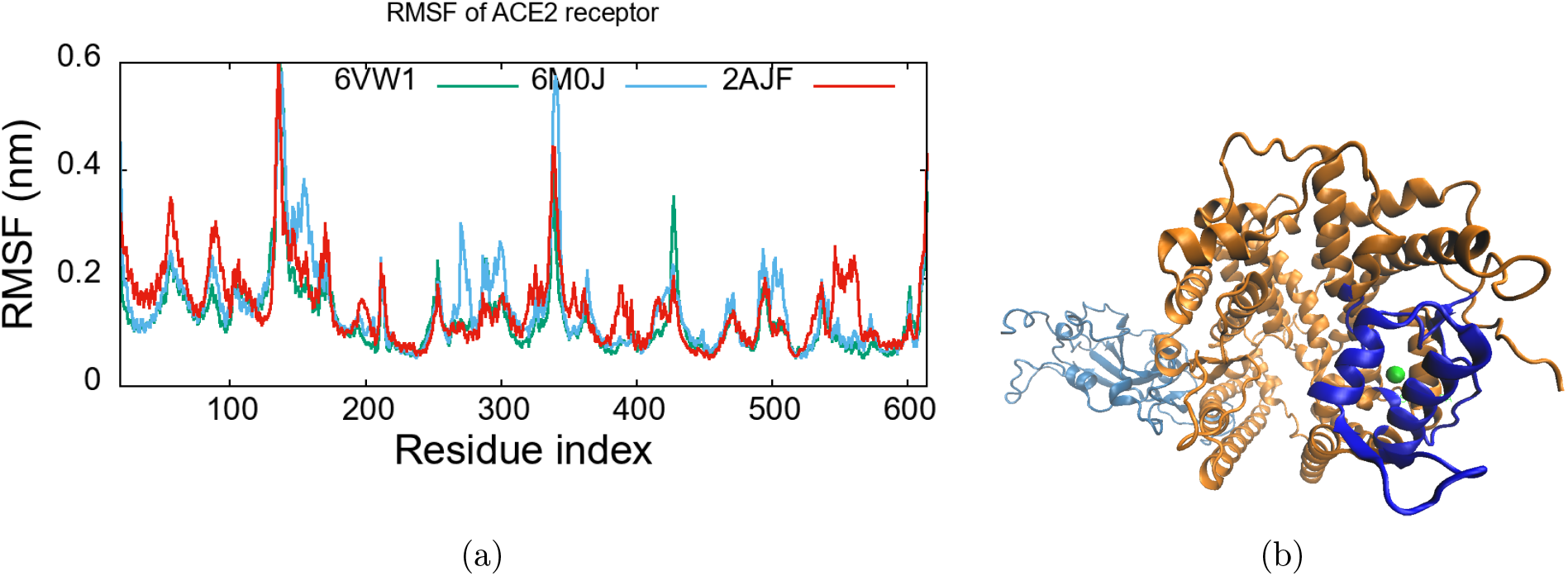
Root-mean-square fluctuations of the backbone C_*α*_ atoms of the human ACE2 receptor as function of the residues along the protein backbone (a) for different viruses. Although very similar to each other, the 6M0J system shows slightly higher fluctuations at certain ranges of residues. Figure (b) show these residue ranges in blue color. They are all located near a Cl^−^ ion (shown in green). See text for more details.

The RMSF for the viral RBD core region is plotted in Fig. 7. Here, instead of following exact residue order, the residue indices are shifted and aligned according to the alignment table shown in Fig. 2. This is done so that similar amino acids of the two viruses are compared to each other. One can see from this figure that despite variations in the exact sequences of the two SARS-CoV-2 samples, they behave very similarly to each other (the blue and green curves). Both of them are more stable, with less thermal fluctuations than the SARS-CoV virus (the red curve). The downward arrows in this figure show the locations of the four mutations highlighted in Fig. 2. The mutation, -PP to GVE, leads to the strongest, most notable difference between the two viruses, with the new coronavirus showing much less fluctuations. This is in line with our earlier suggestion that insertion of Glycine and substitution of rigid Proline make the backbone more flexible, allowing it to move closer to the receptor and to bind tighter to it.

**Figure 7:**
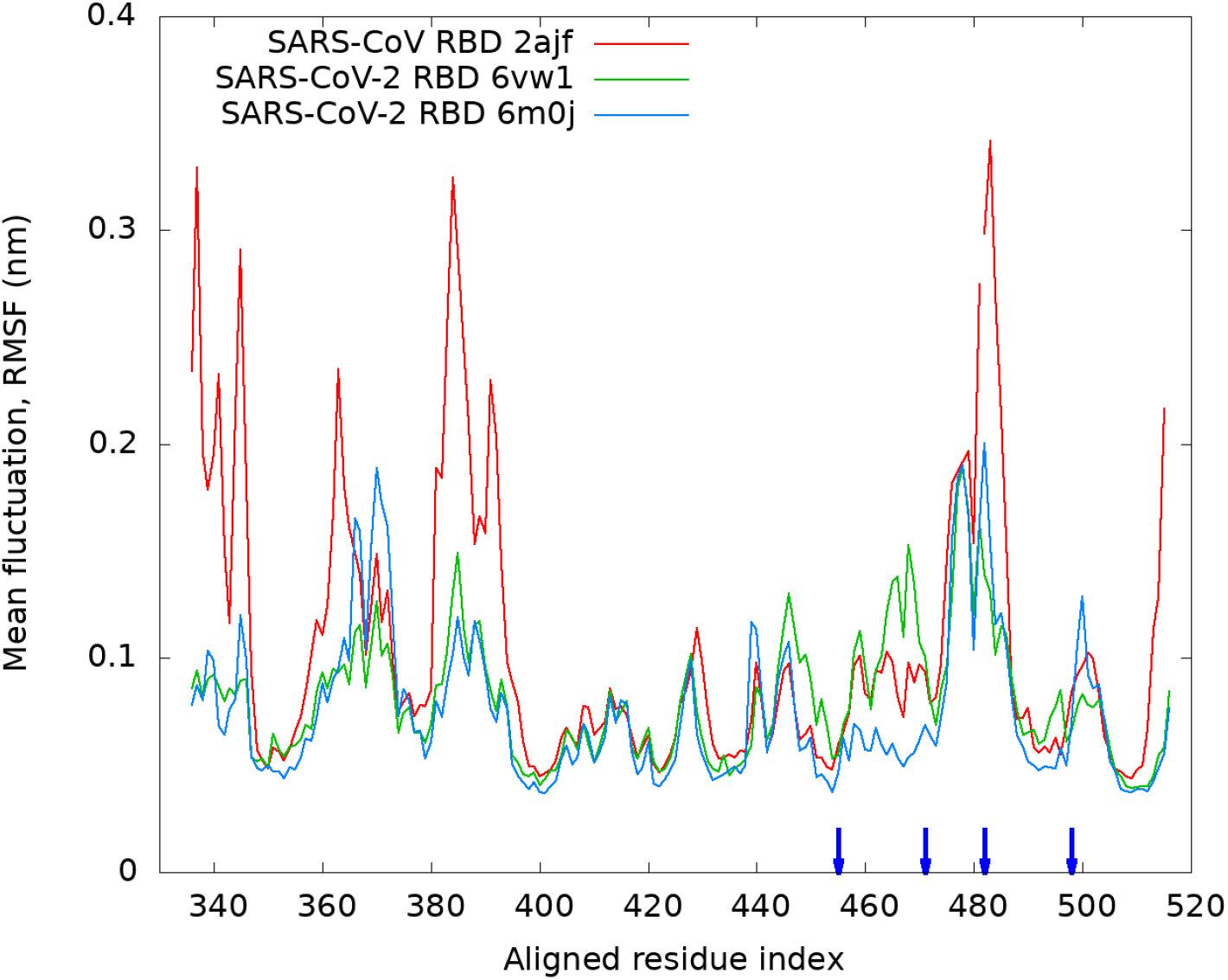
Root-mean-square fluctuations of the backbone C_*α*_ atoms of the viral RBD as function of the residues along the protein backbone for different viruses. Only the core binding region is shown. The residues are aligned and shifted according to Fig. 2 for better comparison between the two viruses.

Remarkably, Figure 7 also shows that, at the N-terminal part, around the indices 340, 365, and 390, the SARS-CoV sample is much less stable than the two SARS-CoV-2 samples. Close inspection of the secondary structure in this region (not shown) show that the various *β*–sheets here are longer in SARS-CoV-2 viruses than those of SARS-CoV virus, indicating a more ordered and compact structure. Stronger receptor binding seems to stabilize not only the ACE2-RBD binding interface (see the next Section), but also the overall structure of viral RBD of the new viruses.

### Hydrogen bonding and coordination number between the viral RBD and the human ACE2 receptor

After performing several sequence and structural investigations, let us move to the interaction picture among amino acids at the binding interface of the viral RBD and its human ACE2 receptor. In Fig. 8(a), the distribution of the number of hydrogen bonds between the viral RBD and the receptor is shown for the three samples. As one can see, both variations of the new SARS-CoV-2 viruses show loss of 2-3 hydrogen bonds compared to the SARS-CoV virus sample. When plotting (not shown) the hydrogen bond distribution for the specific mutated residues highlighted in Fig. 2, it is observed that the first mutation from Tyrosine to Leucine turns off hydrogen bonds at this location due to the lack of side chain hydroxyl (−OH) group in Leucine as expected.

**Figure 8:**
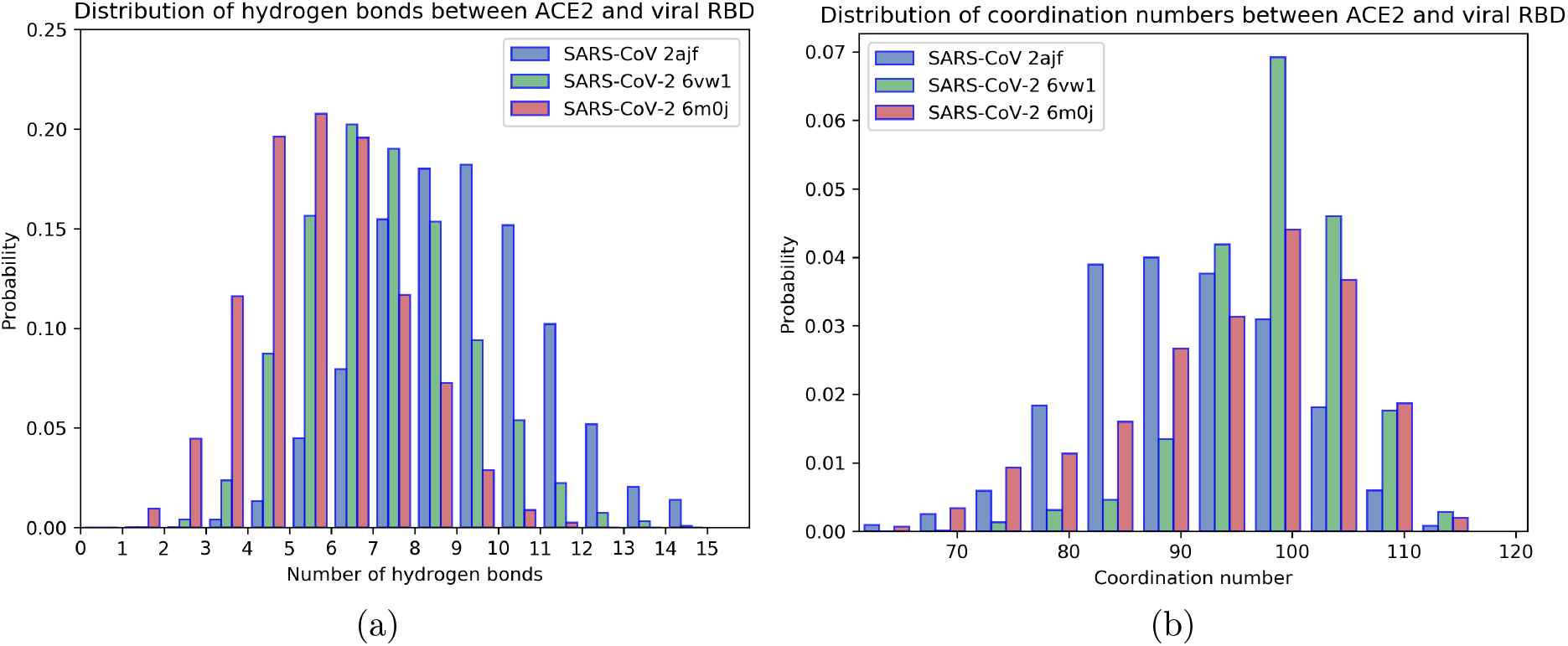
a) Distribution of the number of hydrogen bonds between the viral RBD protein and its human ACE2 receptor. Both SARS-CoV-2 viruses have less hydrogen bonds than that of SARS-CoV virus. b) Distribution of the coordination number between the viral RBD protein and its human ACE2 receptor. Both SARS-CoV-2 viruses have higher coordination number than that of SARS-CoV virus.

It should be noted that the number of hydrogen bonds can only be accurately determined from a full quantum mechanical calculation. However, such calculations are beyond our computing capacity for such a large system. Here, simple geometric criteria are used to determine the hydrogen bonding, namely the distance between the donor and acceptor atoms is less than 3.5 Å, and the angle of the three atoms making up the hydrogen bond is less than 30°. This may overestimate the total number of hydrogen bonds since one atom can participate in more hydrogen bonds at once. Nevertheless, even with this classical definition, the loss of one hydrogen bond due to this mutation is quite clear in our figure.

In addition, in Fig. 8(b), the coordination number between the C_*α*_ atoms of the viral RBD and its receptor is plotted for the three systems in order to estimate the number of interactions between the studied RBDs and ACE2 receptor. Here, the PLUMED driver version 2.5.1^43^ is used for the calculation, for which the default contact function for a pairs of atoms *i* and *j* is given by:

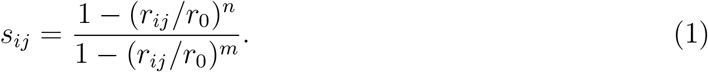

Here, *r_ij_* is the distance between the atoms, *r*_0_ is the neighbor cutoff distance (set to 8Å in our calculation), *n* = 6, and *m* = 12. The meaning of the contact function is similar to inverted step function: when *r_ij_* < *r*_0_, *s_ij_* ≃ 1, when *r_ij_* > *r*_0_, *s_ij_* ≃ 0. Therefore, the contact function denotes whether the two atoms *i* and *j* are within distance *r*_0_ from each other. The expression, Eq. (1), makes the contact function analytically smooth at *r*_0_.

By summing of the contact function over all possible pairs of C_*α*_ atoms, each from different protein, one gets the coordination number which is a measurement of the number of C_*α*_ atoms of one protein within *r*_0_ distance from those of the other proteins. In other words, it is a good indicator of the neighbour interacting residue pairs between the proteins which covers also non-bonded interactions such as van der Waals, electrostatics or hydrophobic excluded-volume interactions. As one can see clearly from Fig. 8(b), although variations between 6M0J and 6VW1 lead to different ranges of the coordination numbers, the values of the most probable coordination number are the same for the two samples and both are greater than the most probable coordination number for SARS-CoV virus. Thus, despite losing the hydrogen bond related to Y to L455 mutation, the coordination number (number of contacts) between the proteins increases by 20% in the case of the two SARS-CoV-2 samples, 6M0J and 6VW1. This is more than enough to compensate for the loss of hydrogen bonding, and increases the binding of viral RBD to the receptor for the new viruses.

It is interesting to see that variations in the sequences of 6VW1 and 6M0J samples of the new SARS-CoV-2 viruses only lead to differences in the variance (or the width) of the distribution of the coordination number and/or the number of hydrogen bonds. Specifically, the 6M0J sample shows significant probability for low coordination number and also lower number of hydrogen bonds, perhaps lower binding energy compared to 6VW1 sample. However, these variations do not change the most probable values of the distributions. This suggests that sequence variations in these two SARS-CoV-2 virus samples are not too important for the receptor binding, while the sequence variations from 2AJF SARS-CoV viruses matters more. This fact is similar to many analyses done so far that the two new SARS-CoV-2 viruses behave very much alike each other, and both are different from SARS-CoV virus behavior. This justifies our criteria for selection of the four regions of important mutations as highlighted in Fig. 2.

### Principal component analysis and the free energy landscape in collective variables

Principal component analysis is a useful method to analyze dynamics of proteins.^44^ By calculating the covariance matrix among the atoms, and diagonalize the resultant matrix we obtain eigenvectors of dynamical motions of the protein (similar to the concept of normal modes). This analysis is very helpful for biomolecules since the long–time dynamics will be dominated by only a few lowest energy modes. All the strong oscillatory high energy modes are screened out efficiently using this analysis, leading to a huge reduction in dimensionality of the systems. By our own inspection, three lowest modes are enough to show distinct clusters of configurations for the viral RBD. In Table 4, various information about the dynamics of the proteins using PCA method are listed. Here, the *P_max_*(*a_i_, a_j_*) is the maximum probability density in the plane of the projections (*a_i_,a_j_*) on the principal eigenvectors i-th and *j*-th respectively. The trace of the covariance matrix is a measure of the overall flexibility of the proteins. As one can see from this table, the trace for ACE2 for SARS-CoV and SARS-CoV-2 6VW1 are large and similar to each other. This is expected, since the human ACE2 receptor is a big protein, with same sequence and almost identical native structures in these systems. The ACE2 protein of SARS-CoV-2 6M0J sample shows even a higher value. This may be related to the missing Cl^−^ ion in the experimental structure, and less degrees of glycosylations of ACE2 protein in this sample (see the preliminary Section).

**Table 4:** The trace of the co-variance matrix and the maximum probability density of the projections of the protein backbones on the first three lowest energy modes.

|  | **2AJF** |  | **6VW1** |  | **6M0J** |  |
| --- | --- | --- | --- | --- | --- | --- |
|  | ACE2 | viral RBD | ACE2 | Viral RBD | ACE2 | Viral RBD |
| Trace ($\text{nm}^2$) | 11.719 | 1.797 | 11.943 | 1.216 | 16.649 | 1.220 |
| $P_{\max}(a_1, a_2)$ | 0.263 | 1.133 | 0.137 | 2.476 | 0.169 | 2.616 |
| $P_{\max}(a_1, a_3)$ | 0.231 | 1.148 | 0.154 | 2.338 | 0.159 | 2.782 |
| $P_{\max}(a_2, a_3)$ | 0.309 | 1.607 | 0.228 | 3.744 | 0.274 | 3.152 |

The difference between SARS-CoV and SARS-CoV-2 viruses are apparent from the PCA information for the viral RBD domain. Here, the SARS-CoV 2AJF shows much larger trace of 1.8 nm^2^ while both SARS-CoV-2 samples shows a trace of only 1.2 nm^2^. This is again in line with previous analyses that SARS-CoV-2 viral RBD are more stable.

In Fig. 9, the normalized histogram of the projections of the protein structure on their first two lowest energy modes are shown. The projection on the third mode follows a simple Gaussian distribution (not shown). This map is the probability density of the configuration of the protein in the phase space spanned by these collective variables. Differences in the probability density for the viral RBD are once again significant. The SARS-CoV-2 virus systems show two smooth, Gaussian-like clusters, while that of SARS-CoV shows weak and scattered profiles. The maximum probability densities in Fig. 9 together with projection on the third eigenvector are listed in Table 4 for both ACE2 and viral RBD backbone structures. Using the relation, Δ*G* ~ −*k_B_T* ln*p*, these result suggest that the depth of the free energy landscape of RBD for the SARS-CoV-2 viruses is lower by 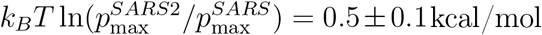. Clearly, the lower fluctuations of the backbone of the viral RBD in the SARS-CoV-2 viruses that we showed in earlier analyses correlate with this lowering in the configurational free energy of the RBD backbone. If one were to contribute this more stable structure due to a stronger binding free energy to the ACE2 receptor, a heuristic estimate for the latter should be about 2-4 times the entropic change, based on the typical balance between interaction energy and entropic configurational freedom for protein-protein complexes.^45^ Our heuristic estimate thus gives a binding free energy difference, ΔΔ*G* of about −1 to −2.5 kcal/mol stronger for the new viruses. Experimentally, it is shown that SARS-CoV-2 virus has 10-20 times higher binding affinity to the receptor than SARS-CoV virus.^27^ Using the relation Δ*G* = *k_B_T* ln *K_d_*, a 10 to 20-fold increase in binding affinity corresponds to ΔΔ*G* in the range of −*k_B_T*ln10 to −*k_B_T*ln20, or about −1.5 to −2.0 kcal/mol. Thus, our heuristic, hand-waving estimate for the binding free energy difference agrees reasonably with this experimental range. One certainly requests a more comprehensive calculation of binding free energy to give more quantitative predictions. However, many traditional semi-empirical methods for quick estimation of free energy which work decently for ligand-protein system fail for protein-protein system because of the large uncertainty related to the huge configurational space of the later. One needs to resort to using various enhanced sampling methods such as metadynamics,^46,47^ umbrella sampling,^48^ or steered molecular dynamics coupled with Jarzynski’s equation.^49^ Different forcefield parametrizations can also lead to variations in this estimate. However, these investigations are extremely resource expensive and are beyond the scope of this work. Here, by many different sequence, structural and dynamical analyses, we have shown that the new SARS-CoV-2 viruses have stronger binding affinity for the human ACE2 receptor than SARS-CoV virus.

**Figure 9:**
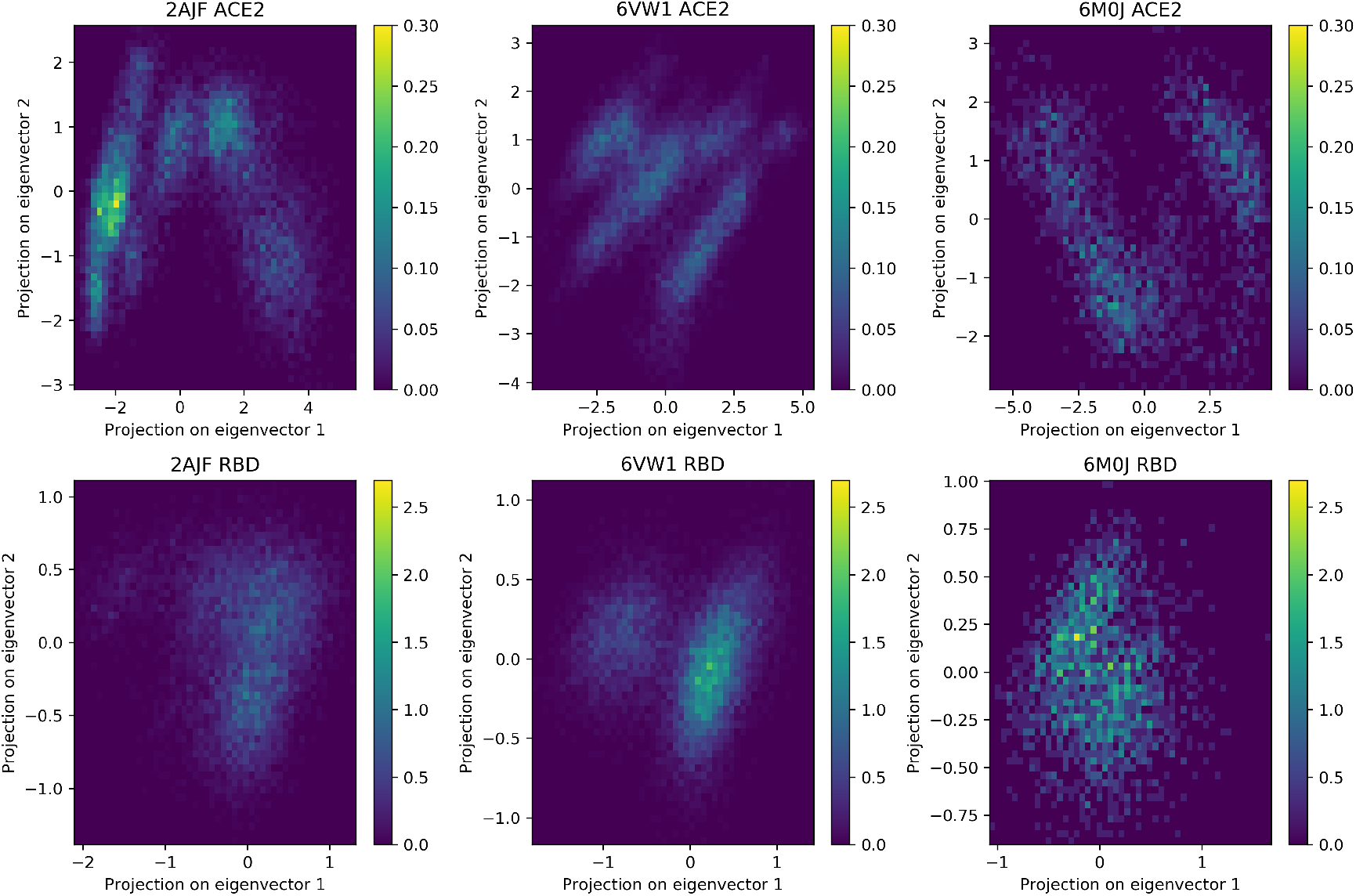
The probability density in the plane of the projections of the structure of the backbones of proteins on its two lowest energy modes. Top row is for the ACE2 receptor and bottom row is for the viral RBD. Note that, the colorbar scale is different for ACE2 versus viral RBD. See text for more discussion.

## Conclusions

In this paper, we present a MD study of the viral receptor-binding domains from SARS-CoV-2 and SARS-CoV viruses in complex with the human ACE2 receptor. By using various sequence, structural and energetic analyses, it is shown that the SARS-CoV-2 viruses bind stronger to its receptor than the SARS-CoV virus. The stronger binding not only creates more stable binding interface, it can also causea more ordered structure of the viral RBD in SARS-CoV-2 virus.

Despite very similar sequence identities, four important mutations in the new viruses play important role in this stronger binding process, namely Y to L455, VP to EI471-472, -PP to GVE482-484, and Y to Q498 (the indices are that of SARS-CoV-2 virus). These mutations all lead to a reduced internal rigidity of the viral protein backbone near its binding interface and add polar and charged residues to the interface. The higher flexibility in the backbone of the new viruses then allows it to move closer to the human receptor protein surface, and to bind stronger to the receptor using non-bonded interactions. Overall, the coordination number between the proteins increases by 20% in the cases of the new SARS-CoV-2 viruses. This increase in different non-bonded interactions (electrostatic, van der Waals, hydrophobic) overcome the loss of one hydrogen bond between the proteins, leading to a stronger binding free energy.

There are some variations in the primary sequences of the two samples of SARS-CoV-2 viruses, with about 85% identity. However, our results show that these variations do not lead to significant differences in the physical properties of the binding complex. Both samples of the new viruses show very similar behaviours. These mutations are therefore non critical for the binding process. Only the non-conserved mutation differences with SARS-CoV are important.

Preliminary estimate of the binding free energy agrees reasonably with the experimental data that SARS-CoV-2 viruses show 10 to 20 folds higher binding affinity for ACE2 receptor as compared to SARS-CoV virus. A more thorough sampling method such as metadynamics, umbrella sampling or steered molecular dynamics is requested for more quantitative comparison. Similarly, the presence of several other mutations in the sequence of new viruses, its variations, stability/conservation are other interesting aspects that needs more thorough investigations. However, all these studies go beyond the current scope of this paper and will be presented in a near future work. Within this work, our sequence, structural and dynamical analyses have provided strong support that SARS-CoV-2 viruses bind stronger to the human ACE2 receptor than SARS-CoV virus.

Our molecular description of the virus-receptor binding interface, particularly the insight we obtained in the role of mutations, can help in the development of vaccines and antiviral drugs by rational drug design addressing the important needs in ongoing pandemics.

## Acknowledgement

TTN acknowledges partial financial support of the Vietnam National Foundation for Science and Technology NAFOSTED grant number 104.99-2016.39. DNM wishes to thank support from the Euratom research and training programme 2019-2020 under grant No.633053 and the RCUK programme under grant No.EP/T012250/1.

We thank Dr. Paolo Carloni and Linh G. Hoang for critical reading of the manuscript.

